# Ergosterol acts as a permissive regulator of Ire1 responsiveness during ER stress

**DOI:** 10.64898/2026.06.19.729409

**Authors:** Porrini Lucía, Almada Juan Cruz, Bortolotti Ana, Antonio D. Uttaro, Germán L. Rosano, Larisa E. Cybulski

**Author notes:** Contributed equally (Porrini Lucía, Almada Juan Cruz). Corresponding autor.

## Abstract

The unfolded protein response (UPR) safeguards endoplasmic reticulum (ER) homeostasis by integrating signals arising from both protein-folding defects and membrane stress. While activation of Ire1 by unfolded proteins has been extensively characterized, the contribution of membrane lipid composition to this process remains incompletely understood. It is unclear whether ergosterol acts as a primary activating signal for Ire1 or instead modulates the activation threshold and amplitude of the UPR response.

Using β-mercaptoethanol (BME) as a proteotoxic perturbation, we found that this reducing agent exerts opposing effects on the two major inputs that converge on Ire1 signaling. Although BME induces proteotoxic stress, it triggers only a moderate UPR response while simultaneously causing a pronounced reduction in ergosterol biosynthesis, a behavior distinct from that generated by classical ER stressors. Pharmacological, proteomic, genetic, and lipidomic analyses revealed a causal relationship between reduced ergosterol levels and attenuated Ire1 activation.

Importantly, elevated ergosterol levels alone were insufficient to activate the UPR, indicating that sterols do not directly trigger the pathway. Instead, our findings support a model in which ergosterol functions as a permissive determinant of Ire1 responsiveness, tuning the amplitude and gain of UPR signaling in response to ER proteotoxic stress. Moreover, the differential effects of endogenous and exogenous sterol accumulation on Ire1 activation raise the possibility that not only total ergosterol abundance, but also its intracellular distribution and accessibility contribute to amplify Ire1 response. Together, these results identify sterol homeostasis as a key regulator of ER stress signaling and reveal how membrane composition influences the efficiency with which luminal stress information is translated into productive Ire1 activation.

## Introduction

The endoplasmic reticulum occupies a unique position at the intersection of protein, lipid, and membrane homeostasis. As the organelle responsible for the synthesis and maturation of secretory proteins, the production of most cellular lipids, calcium storage, and membrane biogenesis, the ER is uniquely positioned to integrate metabolic and structural cues from the cellular environment. Because these processes are tightly interwoven, perturbations affecting protein folding inevitably influence membrane homeostasis, just as alterations in membrane composition can impact protein maturation. Consequently, the ER functions not only as a biosynthetic compartment but also as a major hub for cellular stress sensing and adaptation.

Newly synthesized secretory and membrane proteins fold within the ER lumen, an oxidizing subcompartment enriched in chaperones, glycosylases, and oxidoreductases that collectively sustain protein maturation. Perturbations in this environment—such as redox imbalance, hypoxia, or protein overproduction—compromise folding efficiency and lead to the accumulation of misfolded species. Eukaryotic cells counteract this burden through ER-associated degradation (ERAD), which eliminates aberrant proteins via retrotranslocation and proteasomal degradation, and through the UPR, a signaling network that adjusts ER capacity to restore homeostasis (Walter and Ron, 2011; Radanović and Ernst, 2021; Wiseman, Mesgarzadeh and Hendershot, 2022).

UPR signaling in metazoan cells is mediated by three parallel branches: Ire1, ATF6, and PERK. In contrast, *Saccharomyces cerevisiae* relies exclusively on Ire1, the most ancient and evolutionarily conserved UPR sensor, present from yeast to humans (Patil and Walter, 2001; Calfon *et al*., 2002). Ire1 is an ER transmembrane protein composed of a luminal stress-sensing domain and a cytosolic catalytic region harboring both kinase and endoribonuclease activities (Tirasophon, Welihinda and Kaufman, 1998; Liu *et al*., 2002) (Fig. 1A). Under basal conditions, the luminal domain associates with the chaperone Kar2, which prevents Ire1 activation. Upon disruption of folding homeostasis, Kar2 dissociates from Ire1 and binds unfolded proteins. Under these conditions, Ire1 can also directly capture unfolded proteins. These events promote dimerization and oligomerization, thereby activating Ire1 endoribonuclease activity (Kimata *et al*., 2004; Pincus *et al*., 2010; Le and Kimata, 2021; Radanović and Ernst, 2021). Active Ire1 catalyzes the unconventional splicing of *HAC1* mRNA, which encodes a transcription factor (Cox and Walter, 1996). Once translated, Hac1 translocates to the nucleus and activates a broad transcriptional program involving approximately 300 genes -nearly 5% of the yeast genome-including genes associated with protein translocation, glycosylation, chaperone function, disulfide bond formation, vesicular trafficking, ER-associated degradation, and lipid metabolism (Fig. 1A, Travers *et al*., 2000; Ron and Walter, 2007).

**Figure 1.**
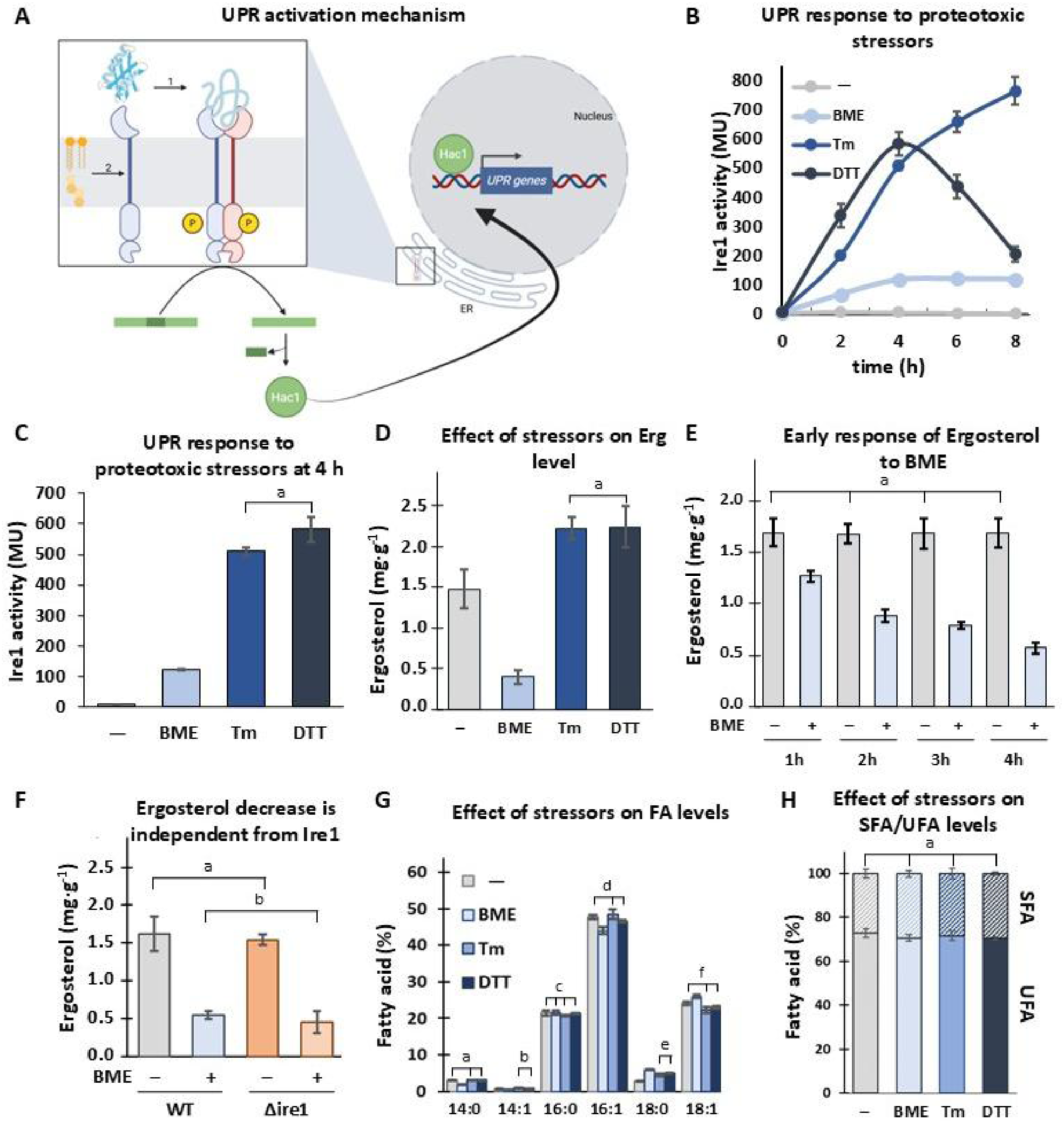
BME induces a moderate UPR while reducing ergosterol levels without altering fatty acid composition. **(A)** Model of Ire1 activation during endoplasmic reticulum stress. Ire1 senses both luminal proteotoxic stress (upper arrow, 1) and changes in the physicochemical properties of the lipid bilayer (lower arrow, 2), triggering oligomerization and Ire1 endoribonuclease activity. *HAC1* mRNA is spliced and translated. Hac1 then is translocated to the nucleus to induce the UPR. **(B)** *S. cerevisiae* cells were transformed with plasmid pJC104 carrying the UPRE–lacZ reporter and were grown in YPD at 30 °C to OD_600_ = 0.4 and then treated with BME (30 mM), Tm (2.5 µg mL⁻¹) or DTT (4 mM). β-galactosidase activity (Miller units) was measured at the indicated time points. **(C)** β-galactosidase activity after 4 h of treatment with the indicated inducers. **(D)** Ergosterol content in cells treated with different stressors was measured by CG-MS. Data are expressed as mg ergosterol per g dry weight of culture. **(E)** Time-dependent decrease in ergosterol levels upon treatment. Ergosterol content was quantified at hourly intervals following treatment with BME. **(F)** Ergosterol content in WT and Δire1 cells treated with BME. **(G)** Fatty acid composition in cells treated with different stressors was quantified by GC-MS. **(H)** Proportion of unsaturated (UFAs) and saturated fatty acids (SFAs) under the indicated conditions. Error bars include the standard deviation from at least three independent experiments. There is no significant difference among values labelled with the same letter.

More recently, evidence has demonstrated that Ire1 also senses alterations in the physicochemical properties of the ER membrane (Promlek *et al*., 2011; Renne and Ernst, 2023; Ernst *et al*., 2024). Perturbations in lipid composition alone are sufficient to activate Ire1. For example, increased levels of saturated fatty acids—either through exogenous supplementation or by deletion of the stearoyl-CoA desaturase—activate the UPR (Volmer, Van Der Ploeg and Ron, 2013). Likewise, alterations in phospholipid composition can induce Ire1 signaling independently of proteotoxic stress (Ho *et al*., 2020; Ishiwata-Kimata, Le and Kimata, 2022). Accumulation of the phospholipid intermediate phosphatidyl-N-monomethylethanolamine (MMPE) triggers chronic ER stress and constitutive Ire1 activation in *S. cerevisiae*, highlighting the sensitivity of the pathway to specific lipid compositional imbalances (Fig. 1A).

Remarkably, an Ire1 variant lacking its luminal unfolded-protein–sensing domain fully retains the ability to detect bilayer stress, demonstrating that Ire1 can directly respond to membrane physicochemical properties (Volmer, Van Der Ploeg and Ron, 2013). Structural studies further proposed that an amphipathic helix adjacent to the transmembrane segment senses lipid packing defects and membrane lateral pressure, providing mechanistic support for the bilayer stress model (Halbleib *et al*., 2017; Väth *et al*., 2021).

Sterols are essential components of eukaryotic membranes and are synthesized in the ER membrane, the same platform where Ire1 resides. Following synthesis, sterols are rapidly distributed to other cellular membranes, maintaining the ER as one of the most sterol-poor compartments in the cell (Daum *et al*., 1998; Mouritsen and Zuckermann, 2004). Ergosterol, the principal sterol in yeast, is tightly regulated through interconnected feedback circuits coordinating sterol biosynthesis, uptake, and membrane homeostasis in response to environmental and metabolic cues (Jordá and Puig, 2020). Thus, although ergosterol is naturally present in the membrane environment surrounding Ire1, the mechanistic relationship between sterol homeostasis and ER stress signaling remains poorly understood.

Beyond their metabolic role, sterols are major determinants of membrane biophysical properties, influencing bilayer thickness, lipid packing, membrane stiffness, and lateral pressure profiles. These parameters critically affect the behavior, partitioning, and oligomerization propensity of transmembrane proteins (Radanović and Ernst, 2021). Consistent with this view, a heme-deficient yeast mutant lacking both sterols and unsaturated fatty acids exhibits constitutive UPR activation, highlighting the intimate connection between membrane lipid composition and ER stress signaling (Pineau *et al*., 2009).

Together, these observations support the idea that ergosterol contributes to shaping the mechanical environment in which Ire1 senses stress. Nevertheless, a key unresolved question is whether ergosterol acts as a primary activating signal for Ire1 or instead functions as a permissive structural determinant that modulates the amplitude of the UPR response.

Here, we examined chemical and genetic perturbations that alter sterol abundance, with particular emphasis on BME. Although BME has been primarily used in higher organisms as an antioxidant and redox-stress modulator with beneficial effects on energy metabolism, oxidative stress, and systemic inflammation (Wong *et al*., 2014), its impact on ER stress signaling has not been systematically characterized. By partially uncoupling proteotoxic stress from sterol homeostasis, BME enabled us to dissect how membrane composition influences the integration of proteotoxic and bilayer-derived inputs into Ire1 signaling.

## Results

### BME uncouples ergosterol homeostasis from maximal UPR activation

We examined the response of Ire1 to BME in *S. cerevisiae*, using the widely used transcriptional reporter system UPRE–lacZ, in which the Ire1-specific UPRE promoter element drives expression of β-galactosidase (Cox and Walter, 1996; Zou *et al*., 2022; Mathuranyanon *et al*., 2015).

Reporter activation was measured in BY4741 cells treated with BME and compared with that induced by tunicamycin (Tm) or dithiothreitol (DTT), two well-characterized UPR inducers. Dose–response curves were generated for each compound, including concentrations previously reported in the literature (Cox and Walter, 1996; Halbleib *et al*., 2017) (Fig. S1). Based on these analyses, 30 mM BME was selected for all subsequent experiments, as it produced growth behavior comparable to that observed with 2.5 μg mL⁻¹ Tm or 4 mM DTT, which are commonly used concentrations for these stressors. Under these conditions, BME induced only ∼20% of the UPR activation observed with Tm or DTT. Even at concentrations up to tenfold higher than those used for DTT, BME failed to reach the levels of activation induced by the classical proteotoxic stressors (Fig. 1B–C and Fig. S1).

Given that membrane lipid composition acts as an input for UPR activation, we investigated whether proteotoxic stress reciprocally reshapes membrane lipid composition, focusing on ergosterol levels. Ergosterol content was quantified after 4 h of treatment with each stressor. Remarkably, ergosterol levels decreased by 73% in BME-treated cells compared with untreated controls. In contrast, treatment with either DTT or Tm resulted in an approximately 50% increase in ergosterol content (Fig. 1D, Table SII). This divergent effect of ER stressors on ergosterol levels is striking and, to our knowledge, has not been previously reported. Notably, the increase in ergosterol observed upon DTT and Tm treatment is consistent with previous transcriptomic studies showing that these stressors upregulate *HMG2*, which encodes HMG-CoA reductase, the rate-limiting enzyme of the mevalonate pathway (Travers *et al*., 2000).

To examine the kinetics of sterol depletion, ergosterol levels were measured at hourly intervals following BME treatment. Ergosterol content progressively declined over time, decreasing by 27%, 47%, 68%, and 73% during the first four hours, respectively. These results indicate that ergosterol depletion is a rapid response that precedes the peak of Ire1 activation (Fig. 1E).

To determine whether the decrease in ergosterol levels observed upon BME treatment depends on the UPR sensor Ire1, we compared sterol levels in a Δire1 mutant and the corresponding wild-type (WT) strain. The Δire1 strain behaved similarly to the WT strain, with BME treatment causing a comparable reduction in ergosterol content in both backgrounds (71% in Δire1 and 66% in WT; Fig. 1F). These results indicate that BME-induced sterol depletion occurs independently of Ire1 signaling and therefore does not arise as a secondary consequence of UPR activation. Rather, BME directly perturbs sterol homeostasis through a parallel mechanism.

To determine whether BME induces a broad remodeling of membrane lipids or a more selective sterol-associated effect, we next analyzed fatty acid composition by GC–MS (Fig. 1G). BME treatment induced a modest shift toward longer acyl chains. However, despite these compositional changes at the level of individual lipid species, the overall balance between saturated and unsaturated fatty acids remained largely unchanged under all conditions tested (Fig. 1G–H and Table S1), indicating that ER stress does not trigger broad remodeling of lipid saturation or fatty acid chain length.

Taken together, these results support a model in which the attenuated UPR activation observed upon BME treatment is associated with the marked depletion of ergosterol in the ER membrane.

### BME activates the UPR and inhibits ergosterol biosynthesis

Because the sterol-lowering effect of BME was independent of Ire1, we next sought to identify the mechanism underlying this reduction in ergosterol levels. We therefore performed an unbiased quantitative proteomic analysis of BME-treated and untreated *S. cerevisiae* cultures to determine whether BME alters the abundance of enzymes within the sterol pathway and how this remodeling integrates with UPR signaling. The dataset showed high reproducibility among biological replicates (Fig. S2A), with an average Pearson correlation coefficient of 0.98. Principal component analysis (PCA) revealed a clear separation of samples along PC1 between the two experimental groups, indicating that BME treatment is the major source of variance in the dataset (Fig. S2B).

BME induced differential protein expression across multiple compartments (Fig. 2A), with the ER exhibiting a particularly strong directional bias toward upregulation, consistent with ER homeostasis being a primary site of redox perturbation. Volcano analysis identified 224 proteins significantly altered upon BME treatment (permutation-based FDR ≤ 0.05), pointing to a large proteome remodeling (Fig. 2B). The list of differentially expressed proteins (DEPs) can be found in repository data (File 5 in doi.org/10.57715/UNR/V8RLSD).

**Figure 2.**
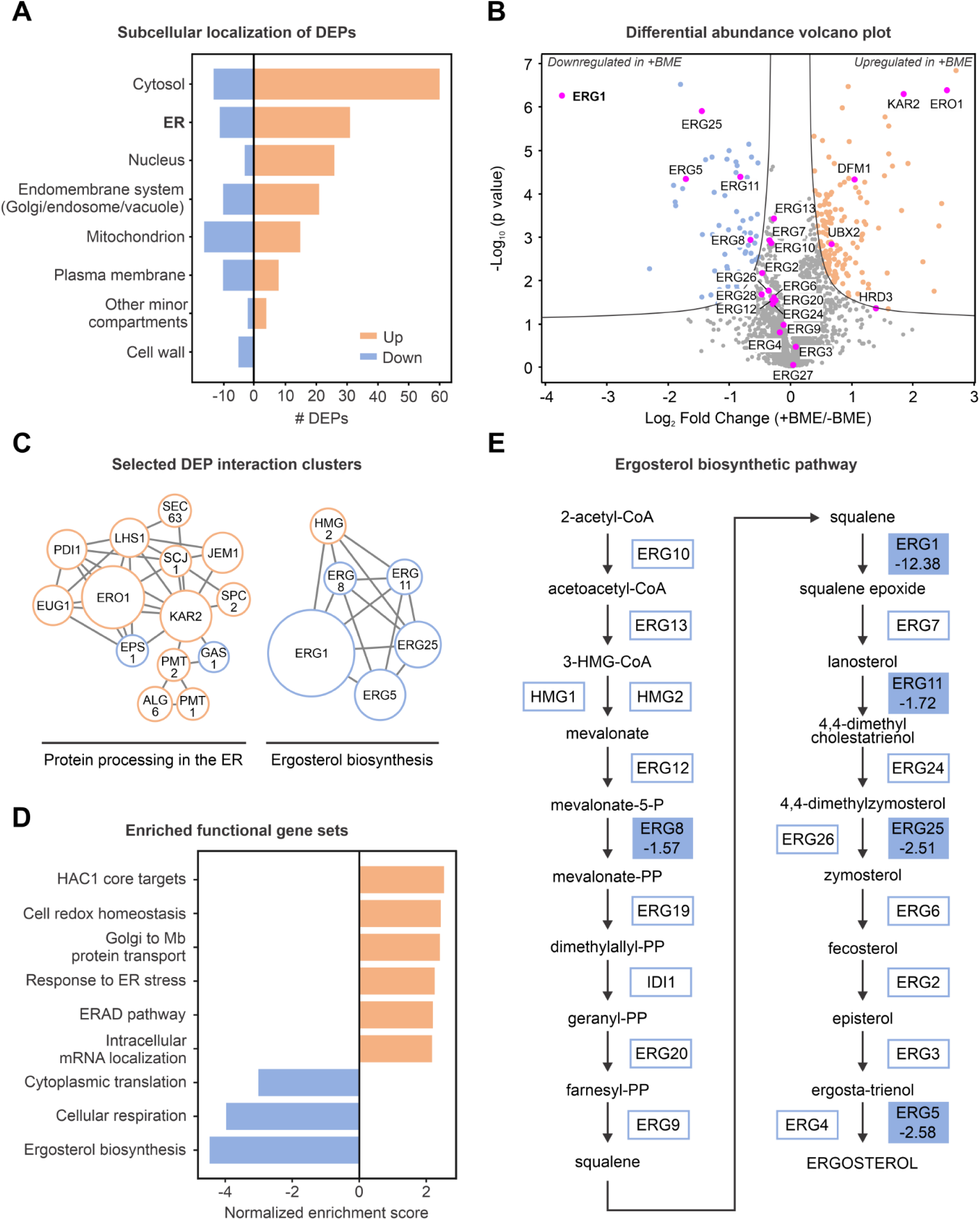
Coordinated repression of the ergosterol biosynthetic pathway and UPR activation upon BME treatment revealed by quantitative proteomics. **(A)** Number of proteins up- or downregulated after BME treatment in different subcellular compartments. The bar plot shows the number of upregulated (orange) and downregulated (blue) proteins across major cellular compartments. **(B)** Volcano plot showing changes in protein abundance in cells treated with BME compared to untreated cells. Several enzymes of the ergosterol biosynthetic pathway, including ERG1, ERG5, ERG11, and ERG25, were among the most significantly downregulated proteins and are highlighted in purple, while signature proteins functioning during protein folding stress in the ER showed the opposite trend. **(C)** Selected protein clusters of DEPs. Node border color indicates down- (blue) or upregulation (orange), while node size reflects fold change. **(D)** GSEA summary plot showing selected enriched gene sets for the +BME condition. Bars represent the normalized enrichment score (NES) for each gene set; positive NES values (orange) and negative NES values (blue) indicate enrichment at opposite ends of the ranked protein list. **(E)** Schematic of the ergosterol biosynthesis pathway. Boxes filled with color denote enzymes identified as differentially expressed in the +BME condition; the numerical value indicates the fold change.

Network analysis revealed two dominant clusters (Fig. 2C): an upregulated protein-processing module enriched in canonical Ire1–Hac1 targets (including Kar2, Pdi1, Ero1, Lhs1, and Eug1, left panel) and a sterol biosynthesis module composed predominantly of downregulated enzymes (Erg1, Erg5, Erg25, right panel). This fact supports a coordinated remodeling of ER proteostasis and lipid metabolism under BME stress conditions (Fig. S3). Because threshold-based approaches capture only strongly changing proteins, we complemented this analysis with Gene Set Enrichment Analysis (GSEA). This rank-based method makes it possible to detect coordinated shifts across predefined biological pathways. This approach confirmed robust activation of the Ire1–Hac1 program, with positive enrichment of ER stress response, redox homeostasis, and ERAD pathways (Fig. 2D, File 6 in doi.org/10.57715/UNR/V8RLSD). In contrast, the ergosterol biosynthetic pathway showed strong negative enrichment, together with decreases in cytoplasmic translation and respiration. Visualization of the ergosterol pathway revealed repression across multiple consecutive enzymatic steps, from Erg1 to downstream lanosterol-derived intermediates, indicating a broad blockade rather than an isolated regulatory effect (Fig. 2E). Importantly, comparison with a recently published quantitative proteomic dataset of yeast treated with Tm and DTT (Platzek *et al*., 2025) revealed substantial overlap in upregulated canonical UPR targets (86 proteins shared across treatments, including Kar2, Pdi1, Ero1, Sil1, Lhs1, Scj1, and Jem1; Fig. S4 and File 7 in doi.org/10.57715/UNR/V8RLSD), confirming that BME robustly activates the conserved Ire1–Hac1 proteostasis module. In contrast, repression of the ergosterol biosynthetic pathway was not significant in the Tm/DTT dataset.

### Reduced ergosterol levels impair full activation of Ire1 in response to canonical ER stressors

Next, we investigated whether the reduction in ergosterol levels induced by BME contributes to the attenuated Ire1 activation observed under this condition. To this end, cells were treated with subinhibitory concentrations of fluconazole—an inhibitor of Erg11—to reduce ergosterol content prior to exposure to proteotoxic stressors. We selected 15 μg/mL fluconazole for subsequent experiments, as this concentration significantly reduced ergosterol levels without affecting cell growth over a 6 h period (Fig. S5). Ergosterol-depleted cells were then challenged with BME, Tm, or DTT following the experimental scheme shown in Fig. 3A.

**Figure 3.**
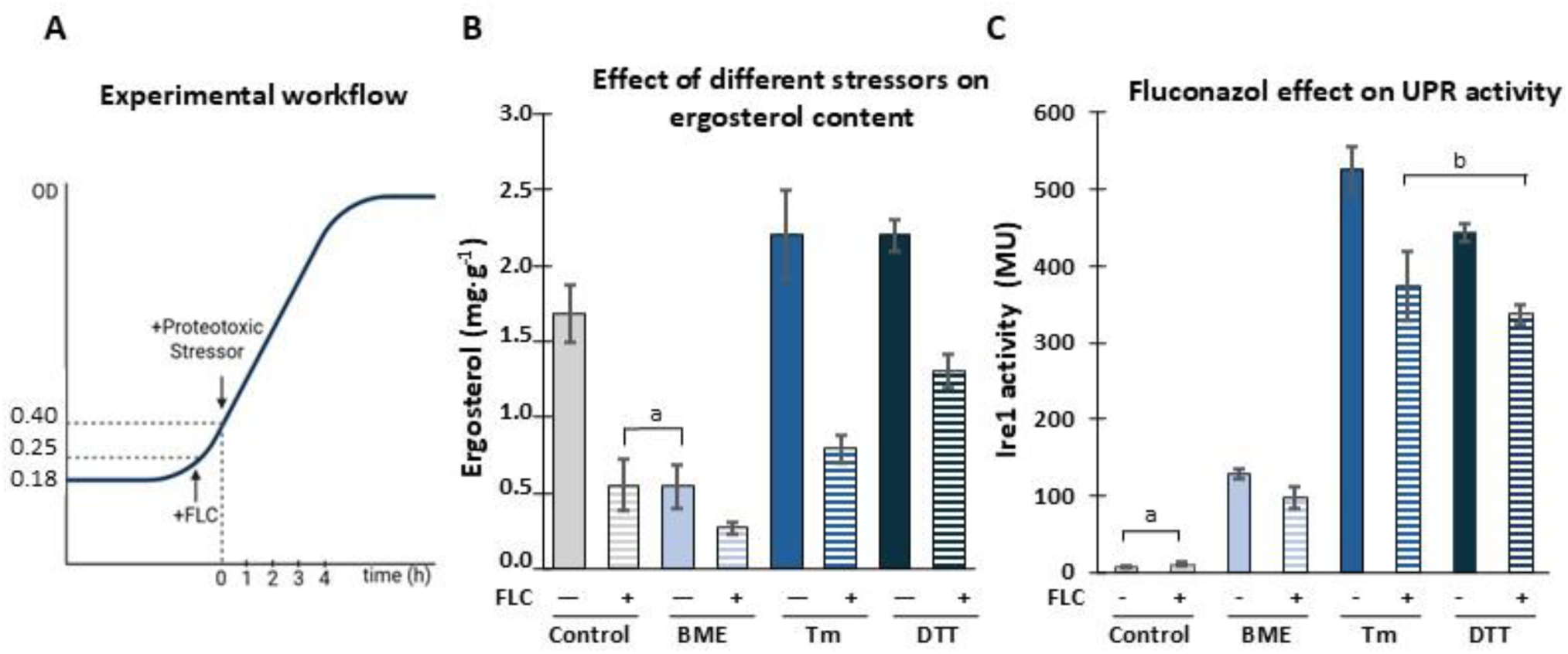
Ergosterol depletion by fluconazole attenuates UPR activation in response to proteotoxic ER stressors. **(A)** Overnight cultures were diluted to OD of 0.18 in synthetic complete medium. When cultures reached an OD of 0.25, cells were treated with or without fluconazole to a final concentration of 15 µg mL⁻¹ (+/-). Upon reaching an OD of 0.4, proteotoxic stressors (BME, TM, or DTT) were added, while control cultures remained untreated. Samples were collected every hour to assess ergosterol content, OD and Ire1 activity. **(B)** Ergosterol levels in *S. cerevisiae* cells treated as described in A. **(C)** Effect of fluconazole on UPR activity. UPR activation was monitored every hour by measuring β-galactosidase activity (Miller units) using the UPRE–lacZ reporter. Bars represent values after 4 hours of treatment. There is no significant difference among values labelled with the same letter. Data of complete curves are available in the repository data (doi.org/10.57715/UNR/V8RLSD). Error bars include the standard deviation from at least three independent experiments.

Cultures treated with either fluconazole or BME exhibited similarly reduced ergosterol levels. The combination of both treatments caused a further decrease, indicating that BME can enhance sterol depletion under conditions of Erg11 inhibition.

A different pattern was observed when fluconazole-treated cells were exposed to Tm or DTT. In these conditions, ergosterol levels were higher than those detected in cells treated with fluconazole alone (Fig. 3B, Table SII). Thus, Tm and DTT partially attenuated the sterol-lowering effect of fluconazole. This effect is consistent with the ability of these stressors to increase ergosterol levels, as observed in Fig. 1D, and suggests that canonical ER stressors can partially compensate for pharmacological inhibition of sterol biosynthesis.

We next evaluated the effect of each stressor, either alone or in combination with fluconazole, on Ire1 activation (Fig. 3C). Fluconazole treatment significantly attenuated UPR activation induced by Tm (∼29%), DTT (∼24%), and BME (∼25%) (Fig. 3C). Notably, sterol depletion reduced Ire1 activation even under the strong proteotoxic stress conditions induced by Tm and DTT. Together these results indicate that unfolded protein accumulation alone is insufficient to sustain maximal UPR signaling in sterol-depleted membranes, supporting a functional link between sterol homeostasis and UPR signaling.

### Strains with elevated endogenous ergosterol levels display an enhanced response to BME Δopi3

To directly discriminate if ergosterol is a signaling trigger that acts synergistically with proteotoxic stressors or is a permissive modulator of Ire1 signaling, we first examined the Δopi3 strain. Opi3 catalyzes the final methylation steps that convert phosphatidylethanolamine (PE) into phosphatidylcholine (PC). Accordingly, Δopi3 cells are deficient in PC synthesis and accumulate PE as well as phosphatidylmonomethylethanolamine (MMPE), resulting in a severe imbalance in phospholipid composition. Previous studies demonstrated that this lipid imbalance is sufficient to induce chronic ER stress and activate Ire1 independently of canonical proteotoxic stimuli (Ishiwata-Kimata, Le and Kimata, 2022). Consistent with these observations, we found that in the absence of proteotoxic stressors Δopi3 cells exhibited markedly elevated levels (107 MU, light green bar, Fig. 4A), whereas the wild-type strain displayed only basal UPR activity.

**Figure 4.**
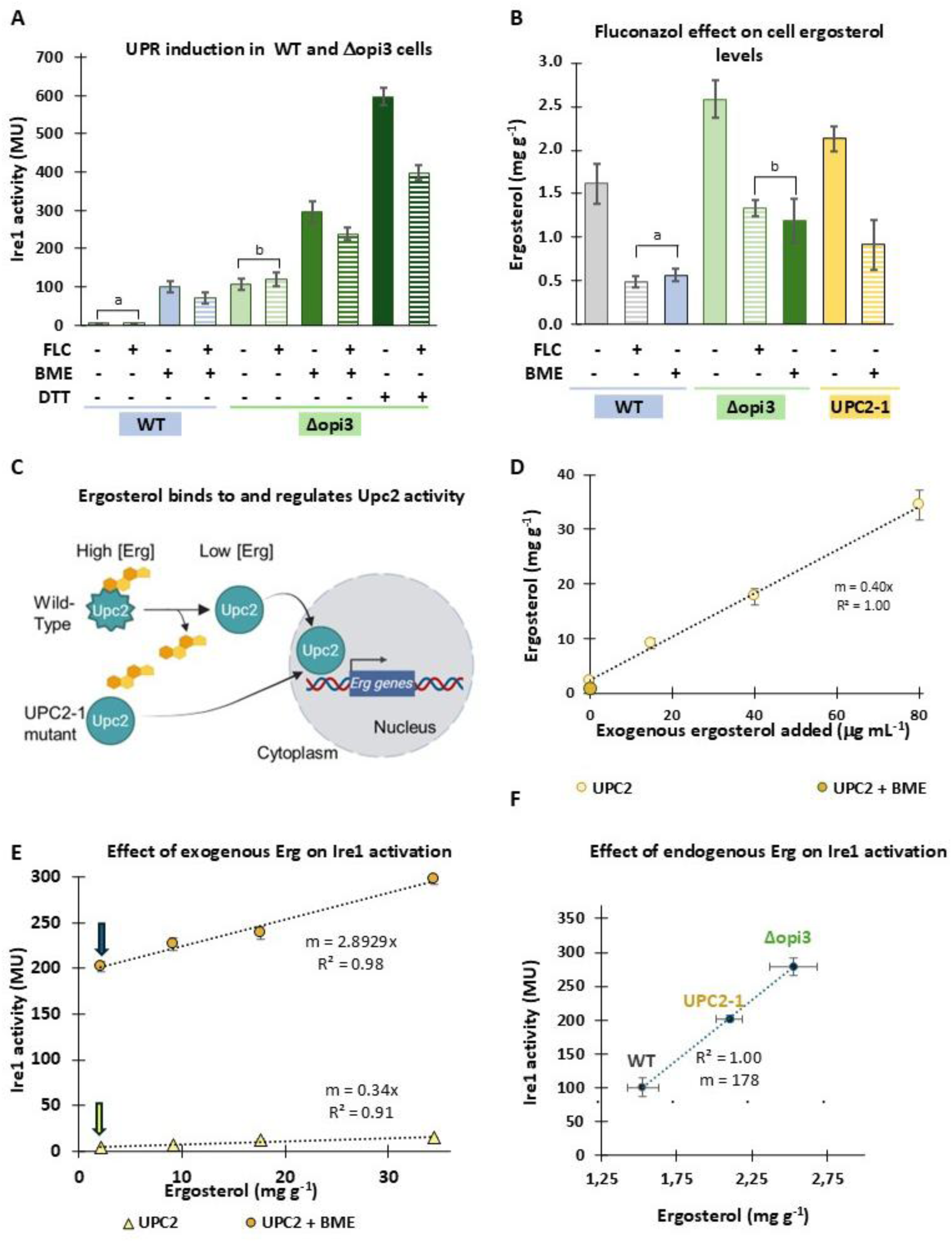
Amplitude of UPR activity correlates with ergosterol levels across distinct genetic backgrounds. **(A)** UPR activity measured using the UPRE–lacZ reporter in wild-type and Δopi3 strains in the presence of fluconazole (FLC) and/or proteotoxic stressors. Values sharing the same letter are not significantly different. **(B)** Ergosterol content in wild-type, Δopi3 and UPC2-1 cells in the presence of FLC and BME. **(C)** Schematic representation of the sterol-responsive transcription factor Upc2. In its sterol-bound state, Upc2 adopts an inactive conformation retained in the cytosol. Upon sterol depletion, ligand dissociation enables nuclear translocation and activation of genes involved in ergosterol synthesis and transport (MacPherson *et al*., 2005) Yang et al., 2020). The G888D mutation disrupts sterol binding and renders Upc2 constitutively active. **(D)** Ergosterol content in the UPC2-1 strain under basal conditions and after supplementation with increasing concentrations of exogenous ergosterol. **(E)** UPR activity in the UPC2-1 strain supplemented with increasing concentrations of exogenous ergosterol. The x-axis indicates the intracellular ergosterol content measured in each condition. Yellow circles represent BME-treated cells and yellow triangles untreated controls. Arrows indicate the coordinate for UPC2-1 activation in the absence of exogenous ergosterol (blue and yellow arrows for activity with and without BME). **(F)** Correlation between endogenous ergosterol levels and UPR activation across different *S. cerevisiae* genetic backgrounds. Each point represents the mean of at least three independent experiments measured 4 h after BME treatment. Error bars stand for the standard deviation.

Because alterations in the PC/PE ratio can trigger compensatory remodeling of other membrane lipids, we reasoned that ergosterol levels might also be altered and participate or be responsible for in Ire1 activation in this strain. Quantification of ergosterol revealed that in fact, Δopi3 cells contained 63% more ergosterol than the wild type (2.6 versus 1.6 mg per gram of dry weight, respectively; green vs grey bars,Fig. 4B). These results suggest that phospholipid imbalance in this strain is accompanied by increased sterol levels, raising the possibility that ergosterol contributes directly to the constitutive activation of Ire1 signaling observed in Δopi3 cells. To determine whether elevated ergosterol is the cause of UPR activation in this mutant, we inhibited sterol biosynthesis using fluconazole. Fluconazole efficiently reduced ergosterol levels in Δopi3 cells (Fig. 4B, striped green bars); nevertheless, Ire1 activity remained strongly elevated even in conditions of ergosterol reduction (119 Miller units; striped green bars Fig. 4A). These results indicate that the phospholipid imbalance present in Δopi3 cells is sufficient to sustain constitutive UPR activation, and that the accompanying increase in ergosterol is not required to maintain this basal activated state. We next asked whether this strain, which contains elevated basal ergosterol levels, could mount an exacerbated response to proteotoxic stress. Upon treatment with BME or DTT, Δopi3 cells exhibited a markedly stronger UPR response than the wild type, increasing increases of 116% and 293%, respectively (Fig. 4A). Interestingly, this enhanced response to BME and DTT was attenuated by fluconazole treatment (striped bars), indicating that elevated endogenous sterol levels in this strain potentiate Ire1 signaling under proteotoxic stress conditions. Together, these results suggest that increased basal ergosterol levels in Δopi3 cells do not directly trigger the UPR, but rather enhance the amplitude of the response once the proteotoxic stress is imposed.

Together, these observations strongly suggest that phospholipid imbalance determines the constitutive activation of the UPR, whereas sterol levels modulate the magnitude of Ire1 signaling during proteotoxic stress.

### UPC2-1

We next used an independent genetic approach to test whether ergosterol acts as a bona fide activating signal for Ire1 in the absence of proteotoxic stress, or instead modulates the sensitivity of the pathway to ER stress. Because wild-type *S. cerevisiae* cells do not import exogenous ergosterol under aerobic conditions, we used the *UPC2-1* mutant strain (Wilcox *et al*., 2002), which enabled controlled modulation of cellular sterol content. Upc2 is a sterol-responsive transcription factor that regulates the expression of ergosterol biosynthetic enzymes and sterol transporters. The *UPC2-1* allele carries the G888D mutation that disrupts sterol binding and renders Upc2 constitutively active (Tan *et al*., 2022). Consequently, UPC2-1 cells overexpress sterol-related genes, including transporters that enable uptake of exogenous ergosterol, a process that is negligible in wild-type cells (Fig. 4C).

We quantified ergosterol content and confirmed that this strain contained higher ergosterol levels than the wild type (2.1 versus 1.6 mg g⁻¹ dry weight; yellow bars, Fig. 4B). Despite its elevated sterol content, this strain did not exhibit constitutive UPR activation in the absence of proteotoxic stressors (Sup Table II).

To test if ergosterol can trigger the UPR in the absence of proteotoxic stressors, we supplemented *UPC2-1* cultures with increasing concentrations of ergosterol (0–80 μg ml⁻¹), within a range previously reported (Pineau *et al*., 2009). After 4 h of growth in ergosterol-enriched medium, cells were harvested, washed, and intracellular ergosterol levels were quantified. Ergosterol incorporation was efficient and increased linearly with increasing extracellular ergosterol concentrations (Fig. 4D). However, elevated ergosterol alone did not trigger the UPR, and activity remained close to basal levels across this range, showing only a minimal increase (Fig. 4E, yellow triangles, m = 0.34). These results indicate that elevated sterol abundance alone is insufficient to trigger robust Ire1 activation. In parallel, aliquots of ergosterol-loaded *UPC2-1* cells mentioned above, were exposed to BME. Under these conditions, UPR activity increased markedly and displayed a strong linear dependence on ergosterol levels: increasing sterol concentrations progressively amplified UPR activation (Fig. 4E, yellow circles, m = 2.89). This enhancement was proportional to intracellular ergosterol abundance, indicating that elevated sterol content potentiates Ire1 activation under proteotoxic stress conditions.

Finally, we integrated the datasets obtained from wild-type, Δopi3, and UPC2-1 strains to examine whether endogenous ergosterol abundance predicts UPR responsiveness across different genetic backgrounds. Despite the distinct metabolic alterations underlying each strain, a striking linear correlation emerged between endogenous cellular ergosterol levels and the magnitude of Ire1 activation following BME treatment (Fig. 4F, m = 178). Notably, this relationship was substantially stronger than that observed upon exogenous ergosterol supplementation in the UPC2-1 strain (Fig. 4E). Together, these results indicate that endogenous sterol abundance is a strong predictor of UPR responsiveness, whereas increases in total cellular sterol levels achieved through exogenous supplementation have a comparatively modest effect on Ire1 activation.

## Discussion

Dysregulation of the UPR is implicated in a broad spectrum of human diseases, including type 2 diabetes, chronic inflammation, and neurodegeneration, underscoring the importance of precisely calibrated ER stress signaling (Lindholm, 2017; Chen *et al*., 2023). While considerable attention has focused on how unfolded proteins activate ER stress sensors, much less is known about how membrane composition shapes the sensitivity and output of these pathways. The distinct cellular response elicited by BME exposure allowed us to identify ergosterol as a critical modulator of Ire1 responsiveness, redefining its role not as a primary activating signal, but as a permissive determinant that tunes the activation threshold and signaling amplitude of the UPR.

A central observation of this study is that BME and DTT, despite both being thiol-reducing agents that interfere with disulfide bond formation, elicited profoundly different effects on sterol homeostasis and UPR signaling. Whereas DTT triggered the canonical adaptive response associated with increased ergosterol synthesis, BME induced a coordinated repression of the ergosterol biosynthetic pathway, leading to marked sterol depletion and only partial Ire1 activation. These differences indicate that the effects of BME cannot be explained solely by reductive stress or impaired protein folding. Instead, BME appears to induce a distinct metabolic and redox state that partially disconnects proteotoxic stress from efficient Ire1 signaling through sterol depletion.

Importantly, ergosterol depletion occurred rapidly following BME exposure and preceded full UPR activation, suggesting that membrane remodeling is not merely a secondary consequence of ER stress, but an early event that conditions subsequent Ire1 responsiveness. Pharmacological inhibition of ergosterol synthesis using fluconazole further supported this model, as sterol depletion attenuated Ire1 activation even in response to classical proteotoxic stressors such as tunicamycin and DTT. In contrast to saturated fatty acids or phospholipid intermediates such as MMPE, which can directly activate Ire1 signaling when accumulated in the ER membrane, elevated ergosterol levels alone were insufficient to strongly activate the UPR (Pineau *et al*., 2009; Ishiwata-Kimata, Le and Kimata, 2022). Collectively, these observations support a model in which sterols do not function as instructive ligands for Ire1 activation but instead create membrane conditions that favor efficient Ire1 signaling. Our results further suggest that sterol localization, rather than total cellular abundance per se, determines the impact of sterols on UPR signaling. This interpretation is consistent with emerging models proposing that sterol function depends not only on abundance but also on subcellular accessibility and distribution across membrane contact networks (Wong, Gatta and Levine, 2019). Exogenous ergosterol supplementation produced only modest effects on basal Ire1 activity despite substantially increasing total cellular sterol levels, whereas endogenous sterol synthesis strongly correlated with UPR responsiveness across multiple genetic backgrounds. These observations raise the possibility that only a specific ER-accessible sterol pool is functionally coupled to Ire1 regulation, while sterols stored in lipid droplets or other membranes remain largely disconnected from Ire1 stress sensing. This distinction may help explain why total cellular sterol abundance is not necessarily predictive of ER stress sensitivity. More generally, these observations suggest that Ire1 responds to the local physical properties of its membrane environment rather than to global cellular sterol abundance.

A plausible mechanistic explanation emerges from the well-established effects of sterols on membrane organization. By increasing bilayer order, thickness, and lateral packing constraints, ergosterol likely reshapes the energetic landscape governing transmembrane helix association and higher-order clustering of Ire1. Rather than triggering activation directly, sterols may lower the energetic barrier required for productive oligomerization once stress has been initiated. This interpretation is highly consistent with emerging models in which membrane-associated stress sensors respond to changes in bilayer material properties rather than to specific lipid species alone. In diverse systems, including the bacterial thermosensor DesK/DesR two-component system (Cybulski *et al*., 2010; Inda *et al*., 2019; Almada *et al*., 2025), the yeast Mga2 pathway (Ballweg and Ernst, 2017), and the PAQR-2 pathway in *C. elegans* (Pilon, 2016; Bodhicharla *et al*., 2018), alterations in membrane rigidity and lipid saturation are actively monitored and coupled to adaptive lipid remodeling programs.

More broadly, this work supports an emerging view of the ER membrane as an active information-processing platform that integrates metabolic state, membrane composition, redox conditions, and proteostasis into unified signaling outputs. By linking sterol homeostasis to the efficiency of ER stress signaling, our study expands current models of lipid bilayer stress and suggests that modulation of membrane material properties may represent a conserved strategy for tuning stress sensitivity across eukaryotic cells.

## Materials and methods

### Yeast Growth Conditions and UPR Reporter Assay

For UPR activity measurements, cells were transformed with plasmid pJC104 carrying the UPRE–lacZ transcriptional reporter (Cox and Walter, 1996). The corresponding *Saccharomyces cerevisiae* cells (see genotype of each strain in table SIII) were grown at 30°C in YPD to mid-exponential phase (OD_600_ = 0.4). Cultures were divided and subjected to the indicated treatments. Samples were collected at 1 h intervals. Cells were harvested by centrifugation, lysed by sonication in Z buffer (60 mM Na_2_HPO_4_, 40 mM NaH_2_PO_4_, 10 mM KCl, 1 mM MgSO_4_, 1 mM β-mercaptoethanol, 0.5 mg/ml lysozyme y 0.3 % Triton X-100 with 5 pulses of 15 seconds at 25 % amplitude in a Q125 sonicator (Q sonica), and β-galactosidase activity was determined as previously described (Cox and Walter, 1996). Activity values were normalized to OD 600nm.

Experiments performed with the Δopi3 strain were conducted in synthetic complete medium, as growth in YPD abolished constitutive UPR activation (Fig. S6), likely due to the presence of choline derivatives in this complex medium that restore phosphatidylcholine synthesis via the Kennedy pathway (Henry, Kohlwein and Carman, 2011).

### Proteomic Sample Preparation and Mass Spectrometry

Overnight yeast cultures grown in minimal medium were diluted to OD_600_ = 0.18 and grown to OD_600_ = 0.4. Cultures were split and either left untreated or treated with stressors for 3 h at 30°C. Cells were harvested, resuspended in PBS supplemented with 1 mM PMSF, and lysed by sonication. After centrifugation, the soluble fraction was retained. Total protein concentration was determined by Bradford assay. For each condition, 30 µg of protein were separated by SDS–PAGE. Gel lanes were fixed, excised, and processed for label-free quantitative proteomics.

Peptide separation was performed on a nanoLC system (Ultimate 3000, Thermo Scientific) coupled to a Q Exactive HF mass spectrometer. Raw files were processed using Proteome Discoverer 2.4 and searched against the *S. cerevisiae* UniProt database. Quantitative analysis was performed using standard LFQ workflows, and downstream statistical analysis was conducted in Perseus.

Proteins above the significance threshold curves in the Perseus Volcano Plot module were considered DEPs (two-sample t-test, 250 randomizations, permutation-based FDR ≤ 0.05, s0 = 0.15). Gene Set Enrichment Analysis (GSEA) was performed on a preranked proteomic dataset derived from log2 protein abundances. Enrichment was tested against GO Biological Process, WikiPathways, and custom ER/UPR-related gene sets. Protein–protein interaction network analysis was performed using the DEPs queried against the STRING database using a minimum interaction confidence score of 0.7. The resulting network was imported into Cytoscape, singletons were removed, and the remaining network was clustered using the Markov Cluster Algorithm. The proteomics dataset was uploaded to PRIDE (accession number PXD075427).

### Lipid Analysis

After 4 h of treatment, 50 mL culture samples were collected and total lipids were extracted according to the method of Bligh and Dyer (50).

### Fatty Acid Quantification

Fatty acid methyl esters (FAMEs) were prepared by transesterification of glycerolipids using sodium methoxide in methanol (Christie and Breckenridge, 1989) and analyzed by gas chromatography–mass spectrometry (GC–MS). Fatty acids were identified by comparison with authentic standards and spectral libraries. Detailed chromatographic conditions are provided in the Supplementary Methods.

### Ergosterol Quantification

After lipid extraction, ergosterol was quantified either by GC–MS or by an enzymatic assay. For GC–MS analysis, lipid extracts were subjected to alkaline saponification followed by hexane extraction, and ergosterol was identified and quantified using authentic standards (Najle et al., 2013). Detailed analytical conditions are provided in the Supplementary Methods. For routine measurements, ergosterol was quantified using a modified enzymatic assay based on the commercial Colestat® kit (Wiener). Lipid extracts were incubated with fungal lipase, sterol oxidase, and peroxidase in the presence of 4-aminophenazone and phenol, generating a quinoneimine dye quantified spectrophotometrically at 490 nm (Robinet *et al*., 2010). Calibration curves were constructed using authentic ergosterol standards (Sigma-Aldrich). Initial experiments included parallel quantification by GC–MS and enzymatic assay to validate concordance between methods (Fig. S7). Following validation, the enzymatic method was used for routine determinations. Ergosterol levels were expressed as mg g⁻¹ cell dry weight.

## Statistical Analysis

All experiments were performed with independent biological replicates as indicated in figure legends. Data are presented as mean ± SEM. Statistical analyses were performed using GraphPad Prism. Comparisons between two groups were conducted using two-tailed unpaired Student’s t-test unless otherwise indicated. For multiple comparisons, one-way ANOVA followed by appropriate post hoc tests was used. A p-value ≤ 0.05 was considered statistically significant.

## Acknowledgements

L.P. is a postdoctoral fellow of CONICET, and J.C.A. was a doctoral fellow from CONICET. A.B., G.L.R., A.D.U., and L.C. are members of the Research Career of the National Scientific and Technical Research Council of Argentina (CONICET).

This work was supported by grants PIP2020-1862 (CONICET) and PICT2020-01034 (ANPCyT) awarded to Larisa Cybulski.

We thank Peter Walter for providing plasmids, Silvia Rossi for providing strains, Diego de Mendoza for critical reading of the manuscript, Pablo Aguilar for helpful discussions, and Monica Hourcade and Guillermo Marcuzzi for assistance with lipid analysis.

## Data availability

The datasets generated during the current study are available in the repository of the Universidad Nacional de Rosario (UNR), https://dataverse.unr.edu.ar/dataset.xhtml?persistentld=doi:10.57715/UNR/V8RLSD and PRIDE (proteomics dataset; accession number PXD075427).

