## Supplementary material for "Ergosterol acts as a permissive regulator of Ire1 responsiveness during ER stress": Suplemental Material

**Supplementary Methods**

**Parameters for GC–MS Fatty Acid Analysis.** Fatty acids were analyzed as previously described (Almada *et al.*, 2025). Briefly, after 4 h of growth under each treatment, 50 mL culture samples were collected and total lipids were extracted according to the method of Bligh and Dyer (50). Fatty acid methyl esters (FAMEs) were prepared by transesterification with 0.5 M sodium methoxide in methanol (Christie and Breckenridge, 1989). FAMEs were then analyzed by gas chromatography–mass spectrometry (GC–MS) using an Agilent 7890B gas chromatograph coupled to an Agilent 5977A mass spectrometer. Samples were injected onto an HP-88 capillary column (100 m × 0.25 mm internal diameter, 0.20 µm film thickness) with helium as the carrier gas at a constant flow rate of 1 mL $min^{-1}$. The injection volume was 1 µL in split mode, and the inlet temperature was maintained at 250 °C. The oven temperature program was as follows: 120 °C for 5 min, increased at 3 °C $min^{-1}$ to 220 °C, and held for 10 min. The mass spectrometer was operated in electron ionization mode at 70 eV, and spectra were acquired in scan mode over an m $z^{-1}$ range of 50–500. Fatty acids were identified by comparing retention times and mass spectra with authenticated standards and the NIST library. Quantification was based on peak area integration and expressed as the percentage of total identified fatty acids.

**Ergosterol Quantification by GC–MS.** Ergosterol was analyzed as previously described (Najle *et al.*, 2013). Briefly, after Bligh and Dyer extraction, the chloroform phase was recovered and evaporated to dryness under a stream of $N_2$. The lipid extracts were then subjected to alkaline saponification: dried samples were resuspended in 1 mL of 2 M KOH in methanol (methanol:water, 1:0.9) and incubated at 80 °C for 20 min protected from light with aluminum foil. Samples were vortexed intermittently during incubation. After cooling to room temperature, 1 mL of distilled water and 1 mL of hexane were added. Samples were vortexed and centrifuged at 1500 rpm for 5 min to allow phase separation. The upper hexane phase was transferred to a clean tube. Extraction was repeated with an additional 1 mL of hexane, and the organic phases were pooled. The combined hexane extracts were concentrated under nitrogen to a final volume of 600 µL and transferred to amber glass vials for GC–MS analysis. Sterol composition was analyzed using a Shimadzu GC-2010 Plus gas chromatograph equipped with an SPB-1 column (30 m × 0.25 mm × 0.25 µm; Supelco). Retention times and mass spectra of detected peaks were compared with those of authentic standards (Sigma-Aldrich) and spectra available in the National Institute of Standards and Technology (NIST) mass spectral library.

**Proteomic Data Acquisition Parameters.** Peptide separation was performed on a nanoHPLC Ultimate 3000 system (Thermo Scientific) using an EASY-Spray ES903 column (50 cm × 50 µm ID, PepMap RSLC C18). The mobile phase consisted of solvent A (0.1% formic acid in water) and solvent B (0.1% formic acid in acetonitrile). The flow rate was 400 nL/min.

The gradient was: 5–35% solvent B over 90 min, 35–90% solvent B over 20 min, followed by 5 min at 90% solvent B.

Mass spectrometry was performed on a Q Exactive HF instrument (Thermo Scientific). Full MS scans were acquired over m/z 300–1800 at a resolution of 70,000 (at m/z 200), with AGC target of 3e6 and maximum injection time of 100 ms. The top 15 most intense precursor ions were selected for MS/MS fragmentation using a normalized collision energy of 27 eV. MS/MS scans were acquired at a resolution of 17,500, AGC target 2e5, maximum injection time 100 ms, and dynamic exclusion of 30 s.

**Proteomic Data Analysis**. Raw data files were processed using Proteome Discoverer 2.4 and searched against the *S. cerevisiae* UniProt database. Precursor and fragment mass tolerances were set to 10 ppm and 0.02 Da, respectively. Carbamidomethylation of cysteine was set as a fixed modification, and methionine oxidation and N-terminal acetylation as variable modifications. Up to two missed cleavages were allowed, and proteins identified by fewer than two unique peptides were excluded from further analysis. For LFQ-based protein quantification, valid intensity values were required in at least three (out of four) samples per group. Remaining missing values were imputed, and DEPs were identified using the Volcano Plot analysis module in Perseus. Protein abundance values were compared using a two-sample t-test with permutation-based false discovery rate (FDR) control. The analysis was performed with 250 randomizations, an FDR threshold of 0.05, and an s0 parameter of 0.15. For preranked GSEA, proteins were ranked using a SAM-style d-statistic calculated as the difference between mean log2 abundance in BME and control samples divided by the pooled standard error plus a fixed fudge factor (s0 = 0.15).

Rankings were ordered in descending d-statistic values. Gene set libraries included GO Biological Process terms, *S. cerevisiae* WikiPathways, and two custom ER-stress-related sets: KEGG protein processing in the endoplasmic reticulum and a conservative HAC1 core UPR target set. Only gene sets containing 10 to 500 members were retained for analysis. Null distributions were generated by size-matched random sampling from the ranked list (1,000 permutations per set size for full-library analysis and 2,000 for targeted re-evaluation of selected gene sets). Normalized enrichment scores (NES) were calculated relative to same-side null distributions, and false discovery rate (FDR) q-values were estimated separately for positively and negatively enriched gene sets using pooled empirical null distributions. Leading-edge subsets were defined as the genes encountered up to the point of maximum ES. All analyses were performed in Python using pandas, numpy, and matplotlib.

**Table SI**. **Fatty acid compositions of total membrane lipid extracts from *S. cerevisiae* BY4741.** Cells were grown in minimal medium at 30 °C to $OD_{600}$ = 0.40 and then treated with BME (30 mM), Tm (2.5 µg $mL^{-1}$) or DTT (4 mM). Cells were harvested after 4h of treatment. Total lipids were extracted and transesterified to yield FA methylesters. The FA methylesters were subjected to gas chromatography-mass spectrometry analysis. Values are the mean of three independent experiments.

% of total fatty acids

| Fatty acid | **CONTROL** | **BME** | **TM** | **DTT** |
|---|---|---|---|---|
| $C_{14:0}$ | 3.1 | 1.8 | 3.2 | 3.2 |
| $C_{14:1}$ | 0.7 | 0.5 | 0.9 | 0.8 |
| $C_{16:0}$ | 21.4 | 21.5 | 20.7 | 21.3 |
| $C_{16:1}$ | 47.8 | 44.0 | 48.5 | 46.6 |
| $C_{18:0}$ | 2.8 | 6.0 | 4.5 | 4.9 |
| $C_{18:1}$ | 24.2 | 26.0 | 22.3 | 23.0 |
| **Total** | 100 | 100 | 100 | 100 |
| **UFAs** | **72.6** | **70.5** | **71.6** | **70.5** |
| **SFAs** | **27.4** | **29.5** | **28.4** | **29.5** |

**Table SII. Ergosterol content and UPR activity under different genetic and treatment conditions.** Ergosterol levels ([Erg], mg $g^{-1}$ dry weight) and Ire1 activity (Miller units, MU) were measured in the indicated strains under basal conditions or after treatment with DTT 4 mM, tunicamycin (Tm) 2.5 μg·$mL^{-1}$, β-mercaptoethanol (BME) 30 mM, and/or fluconazole 15 μg·$mL^{-1}$. ND stands for not determined.

| Strain | Stressor | Fluconazole | [Erg] | Ire1 activity |
|---|---|---|---|---|
| BY4741 | - | - | 1.6 | 7 |
| BY4741 | DTT | - | 2.2 | 583 |
| BY4741 | Tm | - | 2.2 | 510 |
| BY4741 | BME | - | 0.4 | 111 |
| Δire1 | - | - | 1.5 | ND |
| Δire1 | BME | - | 0.5 | ND |
| BY4741 | - | + | 0.4 | 10 |
| BY4741 | DTT | + | 1.3 | 338 |
| BY4741 | Tm | + | 0.8 | 373 |
| BY4741 | BME | + | 0.25 | 97 |
| Δopi3 | - | - | 2,6 | 107 |
| Δopi3 | - | + | 1,3 | 119 |
| Δopi3 | BME | - | 1,2 | 296 |
| Δopi3 | BME | + | ND | 239 |
| Δopi3 | DTT | + | ND | 397 |
| Δopi3 | DTT | - | ND | 596 |
| UPC2-1 | - | - | 2.1 | 4 |
| UPC2-1 | BME | - | 0.9 | 202 |

Table SIII. **Yeast strains used in this study**

| Strain | Genotype | Source |
|---|---|---|
| WT BY4741 | *MATa his3Δ1 leu2Δ0 met15Δ0 ura3Δ0* | (EUROSCARF) |
| Δopi3 | BY4741 *opi3::KanMX4* | (EUROSCARF) |
| UPC2-1 | *MATa ade2-1 can1-100 his3-11,15 leu2-3,112 trp1-1 ura3-1 hap1::Ty1 upc2-1 (G888D)* | (Lewis *et al.*, 1988; Wilcox *et al., 2002*) |
| Δire1 | BY4741 *ire1::KanMX4* | (EUROSCARF) |

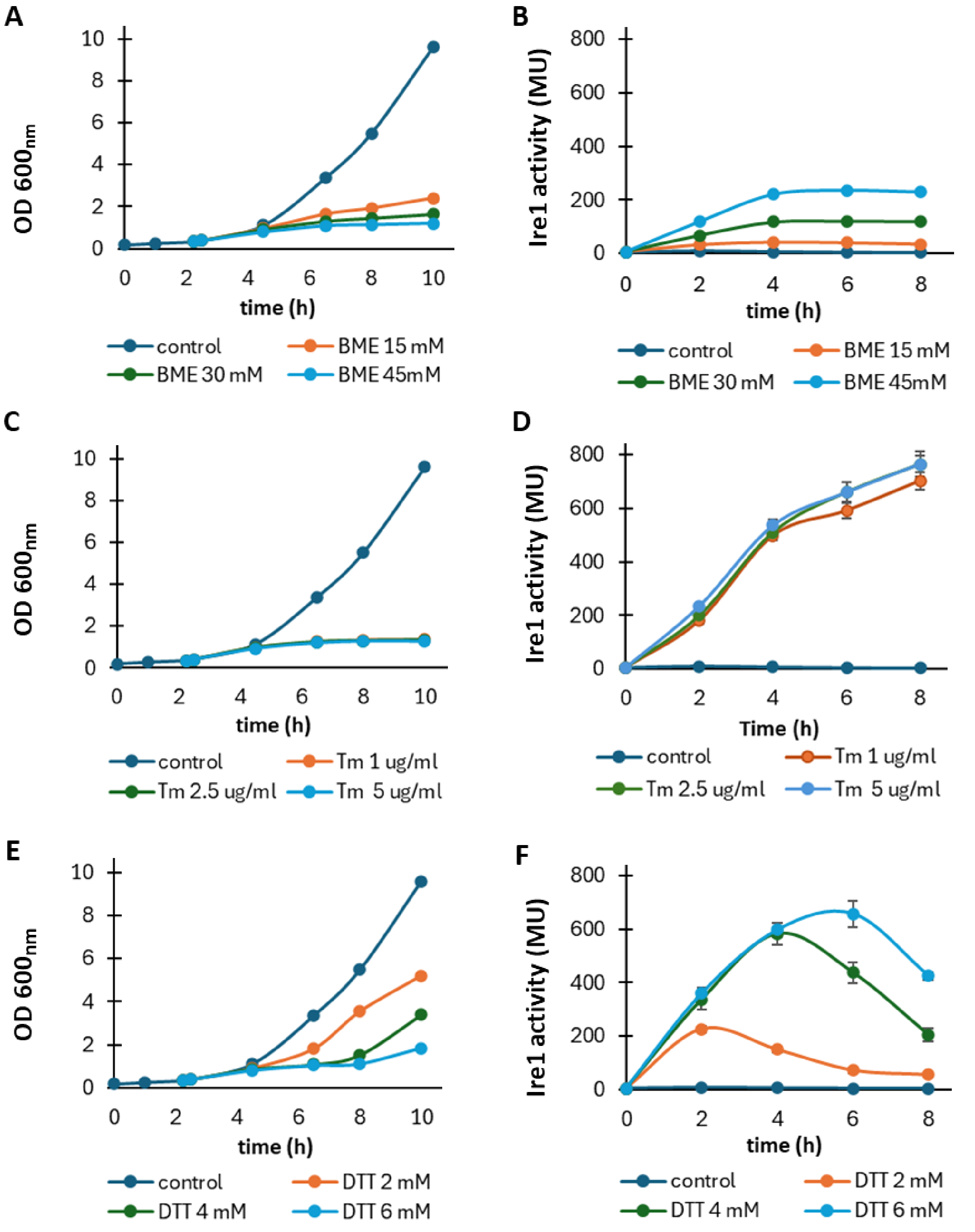


**Figure S1. Growth and UPR activation under different proteotoxic stress conditions. (A, C, E)** Growth curves ($OD_{600}$) of wild-type cells treated at t=0 with increasing concentrations of β-mercaptoethanol (BME), tunicamycin (Tm), or dithiothreitol (DTT), respectively. **(B, D, F)** Corresponding Ire1 activity (Miller units, MU) measured over time using the UPRE–lacZ reporter under the indicated conditions.

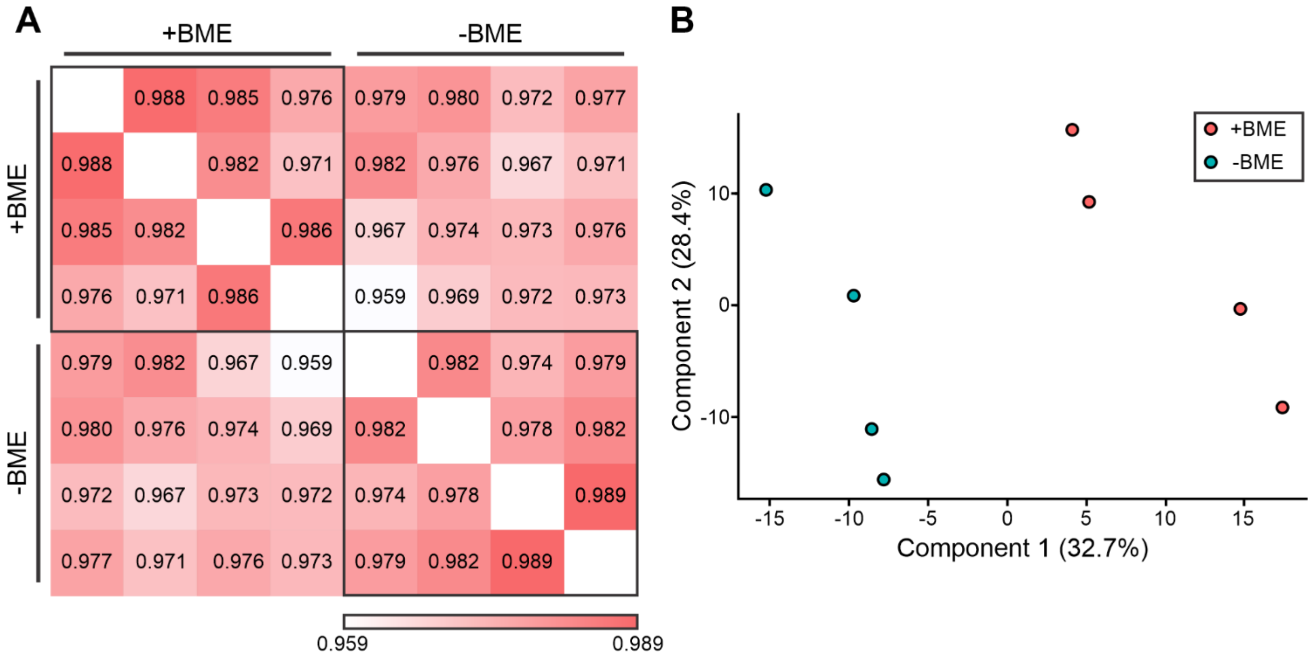


**Figure S2. Quality control and reproducibility of proteomic datasets. (A)** Pairwise correlation matrix of proteomic samples showing high reproducibility across replicates. Color scale indicates correlation coefficients. **(B)** Principal component analysis (PCA) of proteomic samples, showing clustering according to experimental conditions.

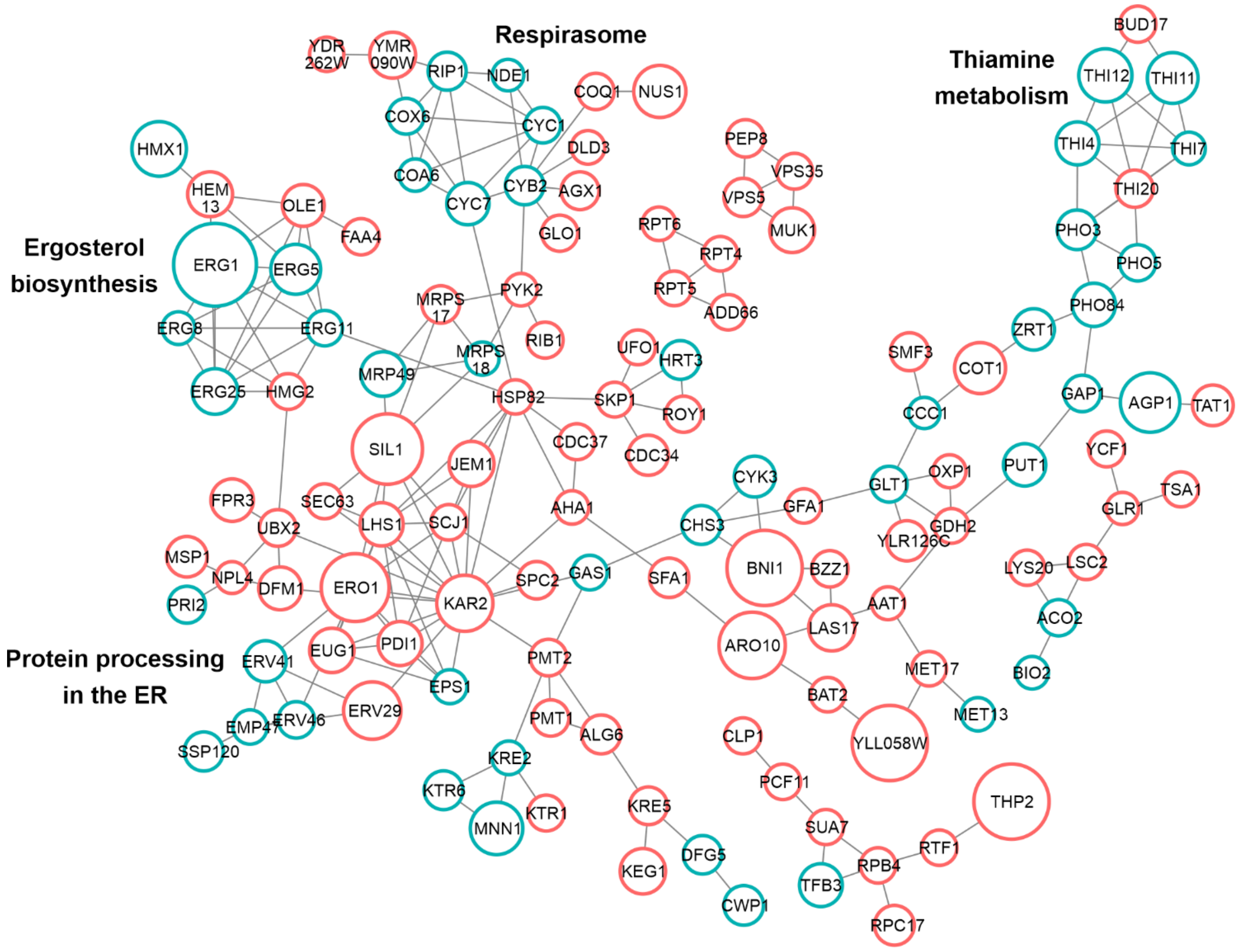


**Figure S3. Protein–protein interaction network of differentially regulated proteins under UPR-inducing conditions.** Node size reflects relative abundance changes, and colors denote differential regulation under the indicated conditions.

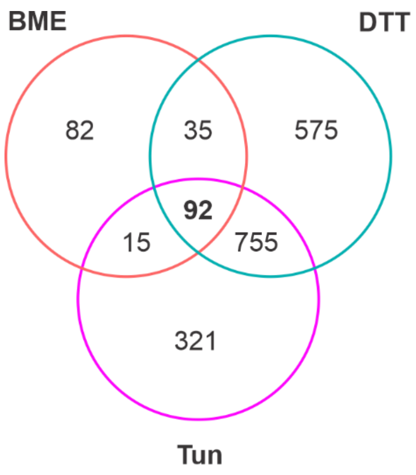

**Figure S4. Overlap between proteomic datasets under UPR-inducing conditions.** Venn diagram showing the overlap of upregulated proteins identified in this study upon BME treatment and those reported in yeast treated with DTT or tunicamycin (Tm) (Platzek *et al.*, 2025). Numbers indicate the count of proteins in each category and intersection.

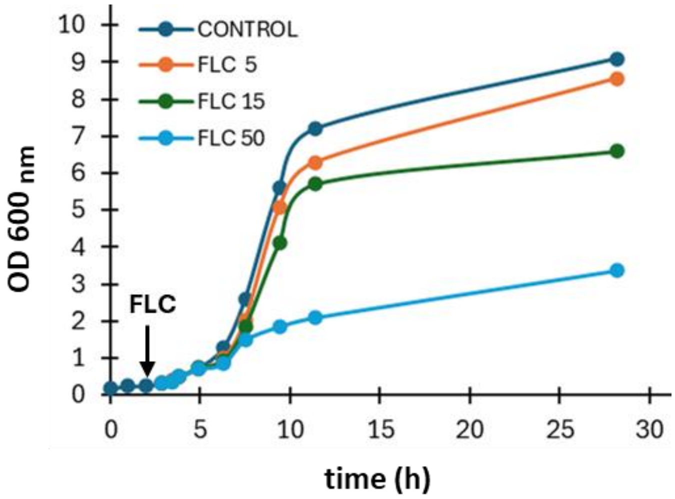


**Figure S5. Growth curves in the presence of increasing fluconazole (FLC) concentrations.** Overnight cultures were diluted to 0.18. After one hour of growth ($OD_{600}$= 0.25), fluconazole was added to reach 5, 15 or 50 µg mL$^{-1}$. OD was monitored for 24 hours. Full datasets are available at doi.org/10.57715/UNR/V8RLSD

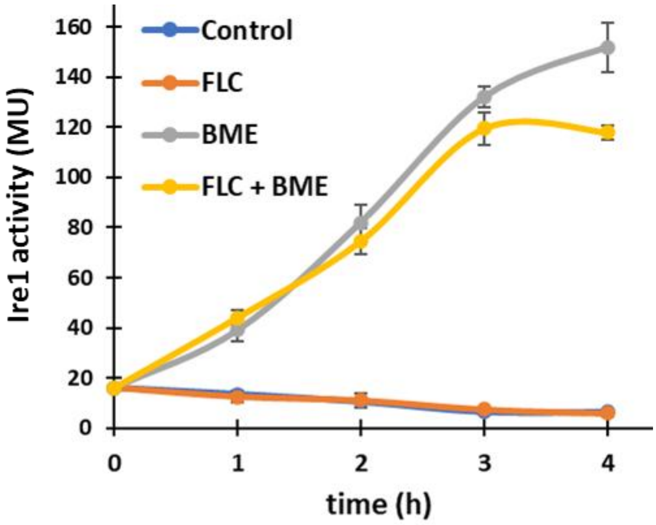

**Figure S6. Ire1 activity in Δopi3 cells in rich medium.** UPR activation was monitored using a UPRE–lacZ reporter (β-galactosidase activity) in Δopi3 cells grown in YPD with or without β-mercaptoethanol (BME) and fluconazole (FLC). Under basal conditions, Δopi3 cells do not display constitutive UPR activation. BME induces a strong time-dependent increase in Ire1 activity, whereas FLC alone has minimal effect. Combined treatment (FLC + BME) partially attenuates BME-induced activation, consistent with sterol depletion modulating the UPR response. Data represent mean ± SD.

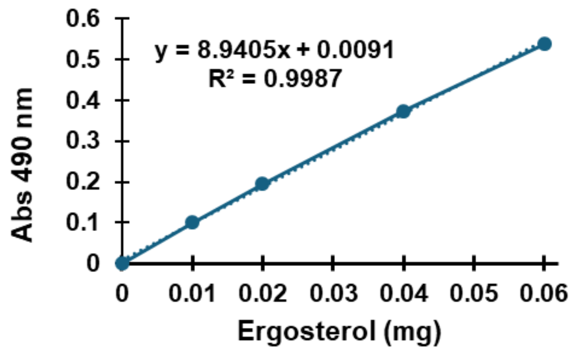


**Figure S7. Ergosterol standard curve for enzymatic assay calibration.** Absorbance at 490 nm was measured for increasing amounts of ergosterol to generate a standard curve. Linear regression analysis showed a strong correlation ($R^2$ = 0.9987), which was used to determine ergosterol concentrations in biological samples.
